# Homologous or Heterologous Booster of Inactivated Vaccine Reduces SARS-CoV-2 Omicron Variant Escape from Neutralizing Antibodies

**DOI:** 10.1101/2021.12.24.474138

**Authors:** Xun Wang, Xiaoyu Zhao, Jieyu Song, Jing Wu, Yuqi Zhu, Minghui Li, Yuchen Cui, Yanjia Chen, Lulu Yang, Jun Liu, Huanzhang Zhu, Shibo Jiang, Pengfei Wang

**Author notes:** These authors contributed equally (Xun Wang, Xiaoyu Zhao, Jieyu Song, Jing Wu, Yuqi Zhu). Correspondence and requests for materials should be addressed to Pengfei Wang.

## Abstract

The massive and rapid transmission of SARS-CoV-2 has led to the emergence of several viral variants of concern (VOCs), with the most recent one, B.1.1.529 (Omicron), which accumulated a large number of spike mutations, raising the specter that this newly identified variant may escape from the currently available vaccines and therapeutic antibodies. Using VSV-based pseudovirus, we found that Omicron variant is markedly resistant to neutralization of sera form convalescents or individuals vaccinated by two doses of inactivated whole-virion vaccines (BBIBP-CorV). However, a homologous inactivated vaccine booster or a heterologous booster with protein subunit vaccine (ZF2001) significantly increased neutralization titers to both WT and Omicron variant. Moreover, at day 14 post the third dose, neutralizing antibody titer reduction for Omicron was less than that for convalescents or individuals who had only two doses of the vaccine, indicating that a homologous or heterologous booster can reduce the Omicron escape from neutralizing. In addition, we tested a panel of 17 SARS-CoV-2 monoclonal antibodies (mAbs). Omicron resists 7 of 8 authorized/approved mAbs, as well as most of the other mAbs targeting distinct epitopes on RBD and NTD. Taken together, our results suggest the urgency to push forward the booster vaccination to combat the emerging SARS-CoV-2 variants.

## Introduction

Coronavirus disease 2019 (COVID-19), caused by severe acute respiratory syndrome coronavirus 2 (SARS-CoV-2), continues to disrupt worldwide social and economic equity, and global public health. As of December 2021, more than 278 million confirmed cases of COVID-19, including 5.4 million deaths, have been reported across the world (https://www.worldometers.info/coronavirus). The massive and rapid transmission of SARS-CoV-2 has led to the emergence of several viral variants, some of which have raised high concern due to their impact on transmissibility, mortality and putative capacity to escape from immune responses generated after infection or vaccination^1^. Distributed throughout the world, the previous four SARS-CoV-2 well-characterized SARS-CoV-2 variants of concern (VOCs), including Alpha (B.1.1.7), Beta (B.1.351), Gamma (P.1), and Delta (B.1.617.2), are responsible for a second and third wave of the pandemic^1,2^. Of note, the Beta (B.1.351) variant displayed the greatest magnitude of immune evasion from both serum neutralizing antibodies and therapeutic monoclonal antibodies, whereas Delta (B.1.617.2) exhibited even greater propensity to spread coupled with a moderate level of antibody resistance^3-6^.

Recently, a new variant of SARS-CoV-2, Omicron (B.1.1.529), was first reported to the World Health Organization (WHO) by South Africa on November 24, 2021. It has been rapidly spreading and was immediately designated as a VOC by WHO within two days^7,8^. Strikingly, analysis of the genomic sequences of the Omicron variant revealed that its spike harbors a high number of mutations, including 15 mutations in the receptor-binding domain (RBD), raising the specter that this newly identified variant may escape from the currently available vaccines and therapeutic antibodies. Moreover, the new variant shares several mutations with the previous VOC Alpha, Beta, Gamma and Delta variants, which further raised global concerns about its transmissibility, pathogenicity, and immune evasion. Here in this study, we constructed the Omicron pseudovirus (PsV) and tested its neutralization sensitivity against convalescent and vaccinee sera as well as a panel of monoclonal antibodies (mAbs).

## Results

A key question in the Omicron investigation thus far is its putative ability to escape immune surveillance. Therefore, we took steps to measure and clarify the extent of such immune evasion by this VOC after infection or vaccination. To accomplish this, we first evaluated the neutralizing activity of serum collected from 10 convalescent patients infected with the Delta variant of SARS-CoV-2 (Supplementary Table 1). Using VSV-based PsVs, we observed robust titers against WT virus in all 10 sera, with the geometric mean neutralizing titers (GMTs) of about 1,100. However, the Omicron variant was markedly resistant to neutralization by convalescent plasma. Only 3 out of the 10 sera showed ID_50_ titers above the lower limit of quantification (LOQ) and a significant drop (>26-fold) compared WT (Figure 1a and Supplementary Figure 1a). We then assessed the neutralizing activity of sera from individuals vaccinated by two doses of inactivated whole-virion vaccines (BBIBP-CorV) (Supplementary Table 1). Here, we found that the GMT against WT was 84 with 80% vaccinees showing positive neutralization activity, while only 10% vaccinees showed successful neutralization against Omicron (Figure 1b and Supplementary Figure 1b). This first impression of Omicron’s immune evasion capability was indeed striking.

**Figure 1.**
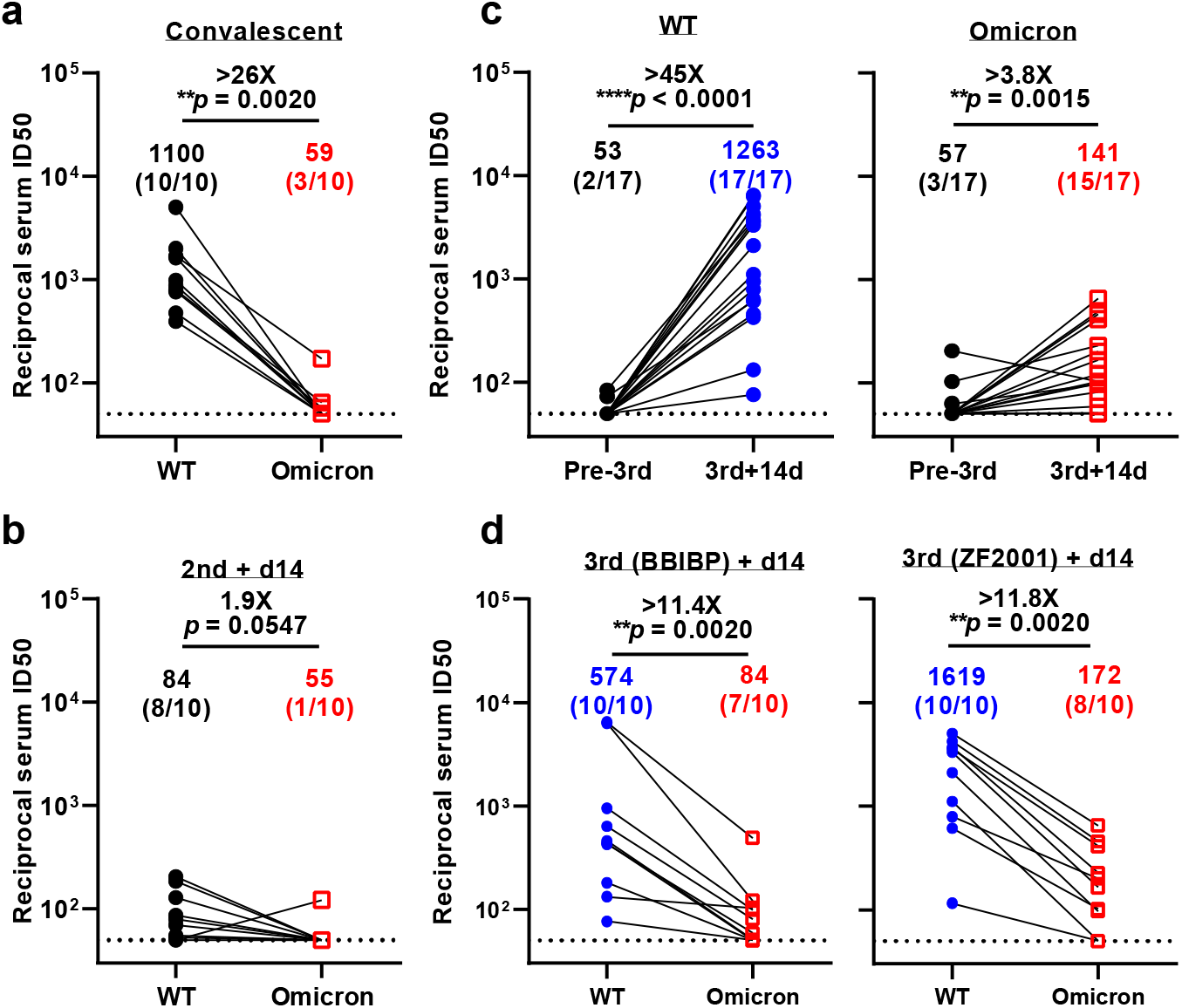
Neutralization of pseudotyped WT (D614G) and Omicron (B.1.1.529) viruses by convalescent sera (**a**), sera collected at day 14 post 2-dose BBIBP-CorV (**b**), sera collected before vs. at day 14 post the booster dose (**c**), and sera collected at day 14 post-BBIBP-CorV or -ZF001 booster dose (**d**). For all panels, values above the symbols denote geometric mean titer and the numbers in parentheses denote the proportion of positive sera with ID_50_ above the LOQ (dotted lines, >1:50). *P* values were determined by using a Wilcoxon matched-pairs signed-rank test (two-tailed).

Then we turned our attention to the efficacy of booster shots now routinely administered in many countries and reported to induce higher immune response^9^. This time, to measure and clarify the efficacy of vaccine boosters against the Omicron variant, we collected samples from healthy adults who had a third boosting vaccination shot with either an inactivated whole-virion vaccine (BBIBP-CorV, homologous booster group) or a protein subunit vaccine (ZF2001, hoterogous booster group) administered at an interval of 4–8 months following previous priming vaccination by two doses of BBIBP-CorV vaccine (Supplementary Table 1). Here, we found that booster vaccine significantly increased the titer of neutralizing antibodies against both the WT and Omicron variant. For the WT virus, the booster improved the neutralization titer more than 45-fold (GMT = 1,263), and for Omicron, the titer increased about 4-fold (GMT = ?, Figure 1c and Supplementary Figure 1c-d). At day 14 post the booster dose, whether homologous or heterologous, the neutralization titer for Omicron was significantly reduced for the both the homologous or heterologous booster, about 11-fold in comparison to WT. However, the reduction fold was less than that for convalescents and the proportion of sera retain active was higher than that of individuals who had only two doses of inactivated vaccine (70-80% *vs* 10%) (Figure 1d and Supplementary Figure 1d). These results indicate that either homologous or heterologous booster can reduce the Omicron escape from neutralizing antibodies.

To understand which types of antibodies in serum lose their activity against Omicron, we further evaluated the neutralization profile of a panel of mAbs targeting SARS-CoV-2 spike (Supplementary Figure 2). These mAbs included most that have been authorized or approved for clinical use: REGN10987 (imdevimab)^10^, REGN10933 (casirivimab)^10^, LY-CoV555 (bamlanivimab)^11^, CB6/LY-CoV016 (etesevimab)^12^, S309 (sotrovimab)^13^, COV2-2130 (cilgavimab)^14^, COV2-2196 (tixagevimab)^14^, and CT-P59 (regdanvimab)^15^, all of which are directed to RBD. Here, we found that 7 out of 8 mAbs tested completely lost their neutralizing activity against the Omicron variant. The only approved antibody that retained its neutralizing activity was S309, but still only about a 7-fold reduction compared to WT (Figure 2, left panel). We further tested some other RBD-directed mAbs of interest, including ADG-2^16^, which is now under development by Adagio Therapeutics, and four more antibodies from our own collection, including 1-20^17^, 2-15^17^, 2-7^17^, and 2-36^17,18^, belong to class 1-4, respectively. The inability of these RBD mAbs to match the risk posed by Omicron was again apparent since all five mAbs tested lost their neutralizing activity, either completely (1-20, 2-15, and 2-7), or partially (ADG-2 and 2-36) (Figure 2, middle panel). We then assessed the neutralizing activity of four N-terminal domain (NTD)-directed mAbs against Omicron and WT viruses, including three targeting the antigenic supersite (5-24, 4-18, and 4-19)^17,19^ and one targeting a distinct site on the NTD (5-7)^17,20^. As shown in the right panel of Figure 2, the activity against Omicron of the supersite-directed mAbs was totally abolished, but only 5-7 retained its activity partially.

**Figure 2.**
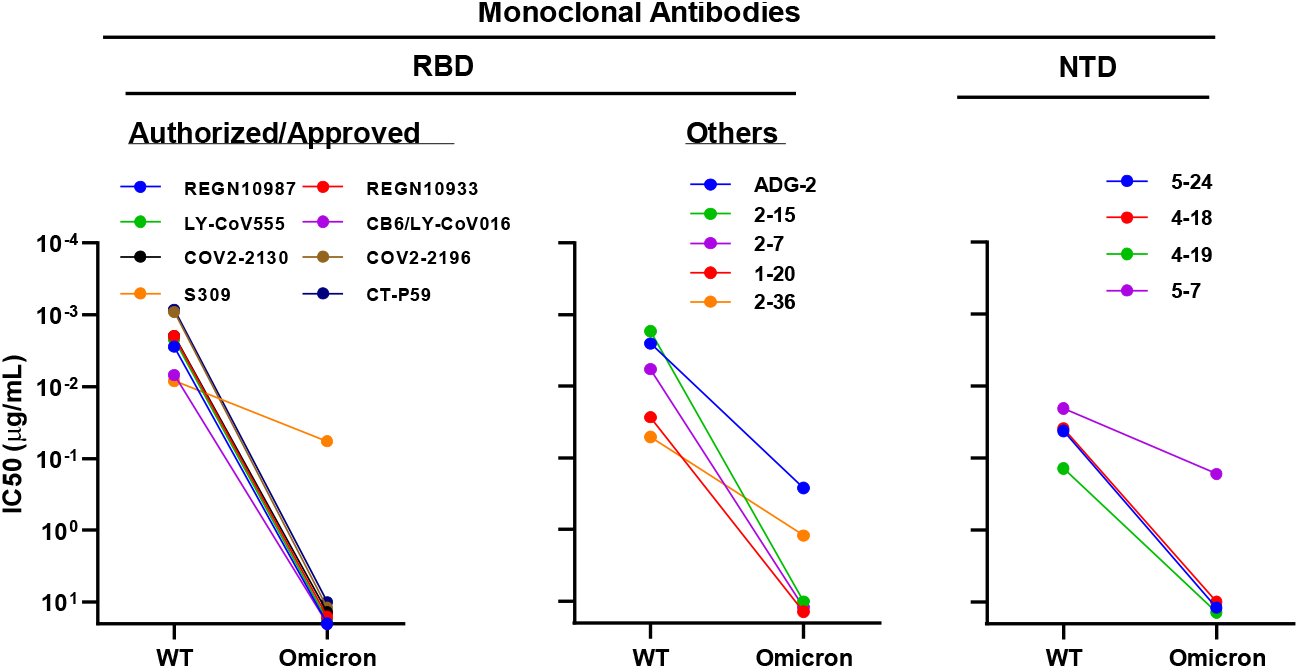
Neutralization of pseudotyped WT (D614G) and Omicron (B.1.1.529) viruses by mAbs targeting different epitopes.

## Discussion

The SARS-CoV-2 Omicron variant struck the world soon after its identification. The retrospective analysis of epidemiological surveillance data in South Africa showed that Omicron is associated with a higher rate of reinfection^21^. Besides, the protection against infection of Omicron from symptomatic disease at 25 weeks after 2-dose vaccination was suggested to be less than 10% in the UK^22^. Similar to these preliminary real-world data, here we reported that a markedly reduced neutralizing activity against Omicron variant in the convalescent and two-dose BBIBP-CorV vaccination group, which is also confirmed by others^23,24^. However, the good news is that after two doses of inactivated vaccines as the “priming” shot, a third homologous inactivated vaccine booster or a heterologous protein subunit vaccine booster could elevate neutralization titer against Omicron. Our results are quite comparable to studies on the same inactivated vaccine^23^ or other vaccines^25^, and again supported by the real-world data showing a third dose booster vaccination provides increased protection against Omicron^26^.

We further investigated the immune evasion capacity of Omicron with mAbs targeting different regions of viral spike protein. Comparable to others^27^, our data suggest the substantial lose of neutralizing activity against Omicron by many mAbs, however, there might be still some conserved sites in RBD (S309 site^13^) or NTD (5-7 site^20^) that could be targeted by antibodies. A vaccine booster, either homologous or heterologous, is expected to elicit such neutralizing antibodies that help reduce the Omicron variant escape and improve the protection. Therefore, it is advisable to push forward booster shots where conditions permit.

## Methods

### Serum samples

Convalescent plasma samples (n=10) were obtained from patients after 3-4 months of SARS-CoV-2 breakthrough infection caused by Delta variant in July 2021. Among them, 9 participants were immunized with two-dose inactivated vaccines (CoronaVac) pre-infection. Sera from individuals who received two or three doses of BBIBP-CorV or ZF2001 vaccine were collected at Huashan Hospital, Fudan University 14 days after the final dose. All collections were conducted according to the guidelines of the Declaration of Helsinki and approved by the Institutional Review Board of the Ethics Committee of Huashan Hospital (2021-041 and 2021-749). All the participants provided written informed consents.

### Monoclonal antibodies

Monoclonal antibodies tested in this study were constructed and produced at Fudan University.

### Construction and production of variant pseudoviruses

Plasmids encoding the WT (D614G) SARS-CoV-2 spike and Omicron variant (B.1.1.529) spike were synthesized. Expi293 cells were grown to 3×10^6^/mL before transfection with the spike gene using Polyethylenimine (Polyscience). Cells were cultured overnight at 37°C with 8% CO_2_ and VSV-G pseudo-typed ΔG-luciferase (G*ΔG-luciferase, Kerafast) was used to infect the cells in DMEM at a multiplicity of infection of 5 for 4 h before washing the cells with 1×DPBS three times. The next day, the transfection supernatant was collected and clarified by centrifugation at 300g for 10 min. Each viral stock was then incubated with 20% I1 hybridoma (anti-VSV-G; ATCC, CRL-2700) supernatant for 1 h at 37 °C to neutralize the contaminating VSV-G pseudotyped ΔG-luciferase virus before measuring titers and making aliquots to be stored at −80 °C.

### Pseudovirus neutralization assays

Neutralization assays were performed by incubating pseudoviruses with serial dilutions of monoclonal antibodies or sera, and scored by the reduction in luciferase gene expression. In brief, Vero E6 cells were seeded in a 96-well plate at a concentration of 2×10^4^ cells per well. Pseudoviruses were incubated the next day with serial dilutions of the test samples in triplicate for 30 min at 37 °C. The mixture was added to cultured cells and incubated for an additional 24 h. The luminescence was measured by Luciferase Assay System (Beyotime). IC_50_ was defined as the dilution at which the relative light units were reduced by 50% compared with the virus control wells (virus + cells) after subtraction of the background in the control groups with cells only. The IC_50_ values were calculated using nonlinear regression in GraphPad Prism.

## Data availability

Materials used in this study will be made available but may require execution of a materials transfer agreement. All the data are provided in the paper or the Supplementary Information.

## Acknowledgements

We are grateful Dr. Rong Zhang (Fudan University) for providing Vero E6 cells. This study was supported by funding from the National Natural Science Foundation of China (82041001 to HZ and 82041025 to SJ).

## Author contributions

P.W. conceived and supervised the project. X.W., X.Z., J.S., J.W., Y.Z. M.L., Y.Cui, Y.Chen, and L.Y. conducted the experiments. J.L. provided materials. X.W., X.Z., J.S., J.W., Y.Z., S.J., and P.W. analyzed the results and wrote the manuscript. All the authors reviewed, commented, and approved the manuscript.

## Competing interests

P.W. is an inventor on patent applications on some of the antibodies described in this manuscript. Others have no conflict of interest.

## References

1 He, X., Hong, W., Pan, X., Lu, G. & Wei, X. SARS-CoV-2 Omicron variant: Characteristics and prevention. MedComm, 1–8, doi:https://doi.org/10.1002/mco2.110 (2021).

2 Davies, N. G. et al. Estimated transmissibility and impact of SARS-CoV-2 lineage B.1.1.7 in England. Science 372, doi:10.1126/science.abg3055 (2021).

3 Mlcochova, P. et al. SARS-CoV-2 B.1.617.2 Delta variant replication and immune evasion. Nature 599, 114–119, doi:10.1038/s41586-021-03944-y (2021).

4 Wang, P. et al. Antibody resistance of SARS-CoV-2 variants B.1.351 and B.1.1.7. Nature 593, 130–135, doi:10.1038/s41586-021-03398-2 (2021).

5 Abu-Raddad, L. J., Chemaitelly, H., Butt, A. A. & National Study Group for, C.-V. Effectiveness of the BNT162b2 Covid-19 Vaccine against the B.1.1.7 and B.1.351 Variants. N Engl J Med 385, 187–189, doi:10.1056/NEJMc2104974 (2021).

6 Liu, C. et al. Reduced neutralization of SARS-CoV-2 B.1.617 by vaccine and convalescent serum. Cell 184, 4220–4236 e4213, doi:10.1016/j.cell.2021.06.020 (2021).

7 Scott, L. et al. Track Omicron’s spread with molecular data. Science 374, 1454–1455, doi:10.1126/science.abn4543 (2021).

8 Callaway, E. Heavily mutated Omicron variant puts scientists on alert. Nature 600, 21, doi:10.1038/d41586-021-03552-w (2021).

9 Ai, J. et al. Recombinant protein subunit vaccine booster following two-dose inactivated vaccines dramatically enhanced anti-RBD responses and neutralizing titers against SARS-CoV-2 and Variants of Concern. Cell Research, doi:10.1038/s41422-021-00590-x (2021).

10 Hansen, J. et al. Studies in humanized mice and convalescent humans yield a SARS-CoV-2 antibody cocktail. Science 369, 1010–1014, doi:10.1126/science.abd0827 (2020).

11 Jones, B. E. et al. The neutralizing antibody, LY-CoV555, protects against SARS-CoV-2 infection in nonhuman primates. Sci Transl Med 13, doi:10.1126/scitranslmed.abf1906 (2021).

12 Shi, R. et al. A human neutralizing antibody targets the receptor-binding site of SARS-CoV-2. Nature 584, 120–124, doi:10.1038/s41586-020-2381-y (2020).

13 Pinto, D. et al. Cross-neutralization of SARS-CoV-2 by a human monoclonal SARS-CoV antibody. Nature 583, 290–295, doi:10.1038/s41586-020-2349-y (2020).

14 Zost, S. J. et al. Potently neutralizing and protective human antibodies against SARS-CoV-2. Nature 584, 443–449, doi:10.1038/s41586-020-2548-6 (2020).

15 Kim, C. et al. A therapeutic neutralizing antibody targeting receptor binding domain of SARS-CoV-2 spike protein. Nature Communications 12, 288, doi:10.1038/s41467-020-20602-5 (2021).

16 Rappazzo, C. G. et al. Broad and potent activity against SARS-like viruses by an engineered human monoclonal antibody. Science 371, 823–829, doi:10.1126/science.abf4830 (2021).

17 Liu, L. et al. Potent neutralizing antibodies against multiple epitopes on SARS-CoV-2 spike. Nature 584, 450–456, doi:10.1038/s41586-020-2571-7 (2020).

18 Wang, P. et al. A monoclonal antibody that neutralizes SARS-CoV-2 variants, SARS-CoV, and other sarbecoviruses. Emerg Microbes Infect, 1–34, doi:10.1080/22221751.2021.2011623 (2021).

19 Cerutti, G. et al. Potent SARS-CoV-2 neutralizing antibodies directed against spike N-terminal domain target a single supersite. Cell Host Microbe 29, 819–833 e817, doi:10.1016/j.chom.2021.03.005 (2021).

20 Cerutti, G. et al. Neutralizing antibody 5-7 defines a distinct site of vulnerability in SARS-CoV-2 spike N-terminal domain. Cell Rep 37, 109928, doi:10.1016/j.celrep.2021.109928 (2021).

21 Pulliam, J. R. C. et al. Increased risk of SARS-CoV-2 reinfection associated with emergence of the Omicron variant in South Africa. medRxiv, 2021.2011.2011.21266068, doi:10.1101/2021.11.11.21266068 (2021).

22 Burki, T. K. Omicron variant and booster COVID-19 vaccines. Lancet Respir Med, doi:10.1016/S2213-2600(21)00559-2 (2021).

23 Ai, J. et al. Omicron variant showed lower neutralizing sensitivity than other SARS-CoV-2 variants to immune sera elicited by vaccines after boost. Emerging Microbes & Infections, 1–24, doi:10.1080/22221751.2021.2022440 (2021).

24 Wang, Y. et al. The significant immune escape of pseudotyped SARS-CoV-2 variant Omicron. Emerging Microbes & Infections 11, 1–5, doi:10.1080/22221751.2021.2017757 (2022).

25 Doria-Rose, N. A. et al. Booster of mRNA-1273 Strengthens SARS-CoV-2 Omicron Neutralization. medRxiv, 2021.2012.2015.21267805, doi:10.1101/2021.12.15.21267805 (2021).

26 Andrews, N. et al. Effectiveness of COVID-19 vaccines against the Omicron (B.1.1.529) variant of concern. medRxiv, 2021.2012.2014.21267615, doi:10.1101/2021.12.14.21267615 (2021).

27 Liu, L. et al. Striking Antibody Evasion Manifested by the Omicron Variant of SARS-CoV-2. Nature, doi:10.1101/2021.12.14.472719 (2021).

