## Supplementary Material for "Homologous or Heterologous Booster of Inactivated Vaccine Reduces SARS-CoV-2 Omicron Variant Escape from Neutralizing Antibodies"

[#]These authors contributed equally

**Supplementary Information and Figures**

**Supplementary Figure 1**. Neutralization curves for convalescent sera (**a**), sera collected at day 14 post the second dose of BBIBP-CorV (**b**), sera collected before the booster dose (**c**), and sera collected at day 14 post the BBIBP-CorV or ZF001 booster dose (**d**).

**Supplementary Figure 2**. Neutralization curves for mAbs.

**Supplementary Table 1.** Bassline characteristics of enrolled participants, including convalescent patients, BBIBP-CorV two doses group, BBIBP-CorV homologous booster group and BBIBP-CorV/ ZF2001 heterologous booster group.

### Supplementary Figures

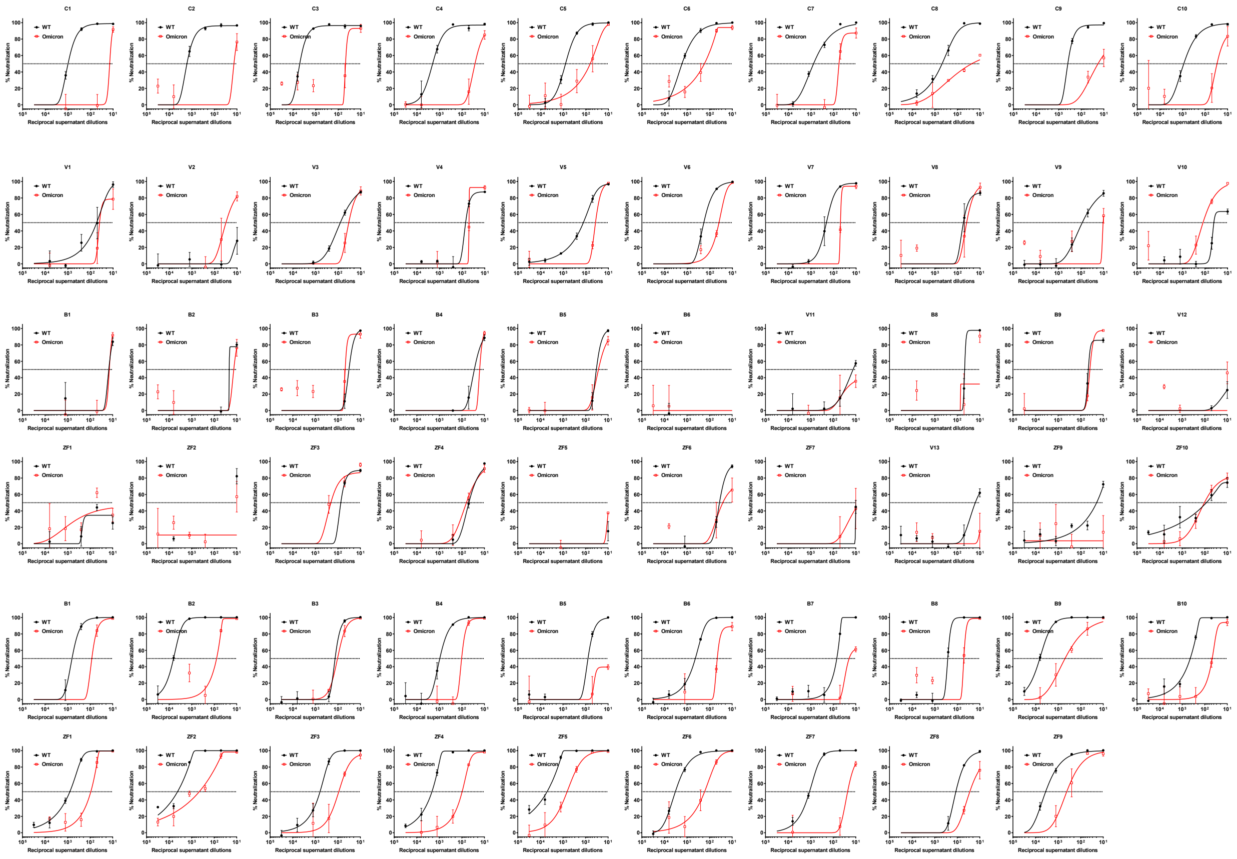


**Supplementary Figure 1.** Neutralization curves for convalescent sera (**a**), sera collected at day 14 post- second dose of BBIBP-CorV (**b**), sera collected before booster dose (**c**), and sera collected at day 14 post the BBIBP-CorV or ZF001 booster dose (**d**).

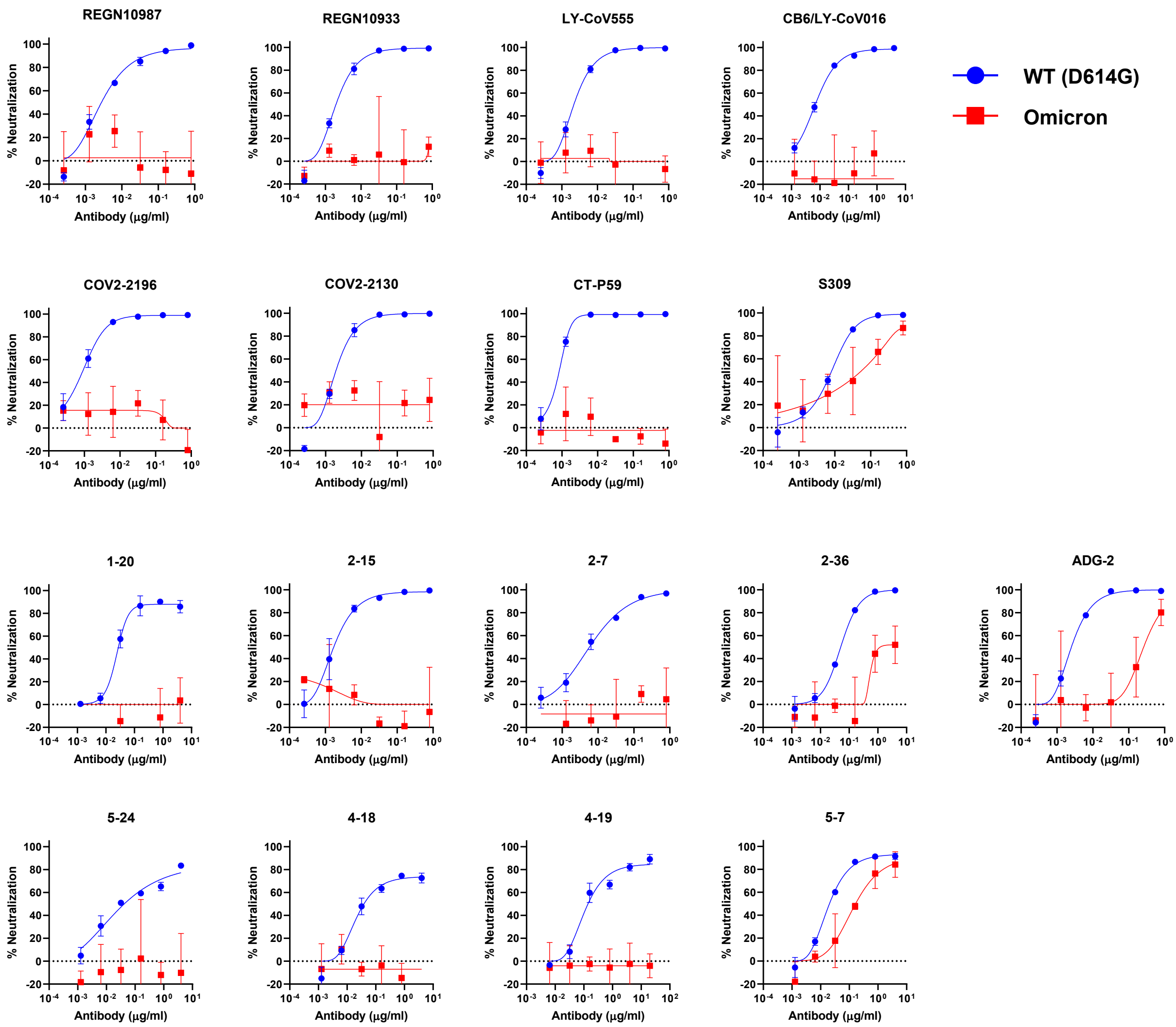


**Supplementary Figure 2. Neutralization curves for mAbs.**

**Supplementary Table 1.** Bassline characteristics of enrolled participants, including convalescent patients, BBIBP-CorV two doses group, BBIBP-CorV homologous booster group and BBIBP-CorV/ZF2001 heterologous booster group.

| | **Convalescent patients** (n=10) | **BBIBP-CorV two doses group** (n=10) | **BBIBP-CorV homologous booster group** (n=10) | **BBIBP-CorV/ ZF2001 heterologous booster group** (n=10) | **P value** |
|---|---|---|---|---|---|
| **Age (years), median(range)** | 46 (34-54) | 33.5 (22-47) | 26(19-31) | 29.5(23-56) | <0.0001 |
| **Male, n (%)** | 3 (30.00%) | 5 (50.00%) | 6 (60.00%) | 45 (50.00%) | 0.684 |
| **BMI (kg/m$^2$), mean (SD)** | 24.45(5.64) | 22.81 (3.27) | 21.61 (3.10) | 21.66 (3.36) | 0.352 |
| **Comorbidities (%)** | | | | | |
| Any, n (%) | 4 (40.00%) | 0 (0.00%) | 0 (0.00%) | 0 (0.00%) | 0.009 |
| Cardiovascular diseases, n (%) | 0 (0.00%) | 0 (0.00%) | 0 (0.00%) | 0 (0.00%) | - |
| Hypertension, n (%) | 3 (30.00%) | 0 (0.00%) | 0 (0.00%) | 0 (0.00%) | 0.049 |
| Diabetes, n (%) | 1 (10.00%) | 0 (0.00%) | 0 (0.00%) | 0 (0.00%) | 1.000 |
